# Estimating the reduction in genetic diversity from background selection under non-equilibrium demography and partial selfing

**DOI:** 10.1101/2025.06.30.662370

**Authors:** Alexander Mackintosh, Maxence Brault, Denis Roze, Martin Lascoux, Sylvain Glémin

## Abstract

The effect of natural selection on linked sites has been suggested to be a major determinant of genetic diversity. While it is in principle possible to estimate this effect from genome sequence data, interactions between selection, demography and inbreeding are expected to make inference less reliable. Here we investigate whether the genome-wide reduction in diversity due to background selection (*B̅*) can be accurately estimated when populations are at demographic non-equilibrium and/or reproduce by partial self-fertilisation. We show that the classic-BGS model is surprisingly robust to both demographic non-equilibrium and low rates of selfing, although both processes do lead to biased estimation of the distribution of fitness effects (DFE) of deleterious mutations. A high rate of selfing leads to poor estimation of both *B̅* and DFE parameters. We propose an alternative inference approach where background selection, demography and partial selfing are jointly estimated from windowed site frequency spectra. This approach, while approximate in nature, resolves most of the bias observed under the classic-BGS model and can also generate estimates of past demography that account for the effect of background selection and partial selfing. We apply the approach to genome sequence data from *Capsella grandiflora* and *C. orientalis*, which have contrasting mating systems and display a forty-fold difference in nucleotide diversity. Our results suggest that background selection has a weak effect on levels of genetic diversity in the outcrosser *C. grandiflora* (*B̅* = 0.89) and a more substantial effect in the predominantly selfing species *C. orientalis* (*B̅* = 0.44), but that background selection alone cannot explain their disparity in genetic diversity.

## Introduction

Genetic diversity varies by several orders of magnitude across species of animals and plants (Leffler *et al*. 2012; Romiguier *et al*. 2014; Chen *et al*. 2017; Buffalo 2021). A number of processes are suggested to contribute to this variation, including differences in *de novo* mutation rate, fluctuations in census population size, population structure and variance in offspring number (reviewed in Charlesworth and Jensen 2022). Although comparative analyses have revealed several correlates of genetic diversity (Romiguier *et al*. 2014; Chen *et al*. 2017), it is still challenging to fully explain differences in diversity between closely related species (Stoffel *et al*. 2018; Mackintosh *et al*. 2019; Barry *et al*. 2022) and we lack a precise explanation for why diversity varies so little in comparison to census population size (Lewontin 1974). The effect of natural selection on linked sites is a potential explanation for at least some of the variation in genetic diversity observed among species. Positive selection acting on a new mutation leaves a surrounding valley of reduced diversity, i.e. a selective sweep, with the width of the valley being inversely proportional to the sojourn time of the selected allele (Smith and Haigh 1974). Recurrent sweeps are expected to have a major impact on genetic diversity in species with large populations and low rates of sexual reproduction / recombination, but to have a much more limited effect in species with small populations and high rates of recombination. Purifying selection also leads to a reduction in genetic diversity at linked sites in a process known as background selection (BGS; Charlesworth *et al*. 1993; Charlesworth 2012). Although selection against a single deleterious mutation has only a very small effect on levels of linked diversity (Hudson and Kaplan 1995; Nordborg *et al*. 1996), deleterious mutations are common enough so that their combined effect can shape the landscape of diversity across genomes and, potentially, lead to considerable differences in genetic diversity between species.

Evaluating the role of BGS in determining levels of genetic diversity within and between species requires estimation of BGS from sequence data. Hudson and Kaplan (1995) and Nordborg *et al*. (1996) derived expectations for the scaled reduction in diversity (*B*) due to purifying selection acting on a deleterious mutation at some recombination distance away. Importantly, they showed that this expectation can be used to predict patterns of nucleotide diversity along the genome, provided that natural selection is strong relative to drift and that the frequencies of deleterious mutations are independent of one another. This model, hereafter referred to as classic-BGS, has since been used to estimate the reduction in diversity due to BGS in model systems such as humans and *Drosophila* (McVicker *et al*. 2009; Comeron 2014; Elyashiv *et al*. 2016; Murphy *et al*. 2022), as well as in a handful of non-model species (Liang *et al*. 2022; Pope *et al*. 2023). Most notably, Corbett-Detig *et al*. (2015) used classic-BGS theory to estimate the combined effect of BGS and recurrent sweeps on genetic diversity across 40 species of animals and plants. They found that species with larger census size experience a greater reduction in genetic diversity from selection at linked sites, and suggested that this effect explains the narrow range of genetic diversity levels observed across species. Although the strength of this conclusion has since been challenged (Coop 2016; Buffalo 2021), comparative analyses like that of Corbett-Detig *et al*. (2015) are an undoubtedly useful approach for understanding the role of selection at linked sites in determining levels of genetic diversity in nature.

Classic-BGS theory assumes that a population is at demographic equilibrium, yet recent work has shown that demographic change modulates the effect of BGS on genetic diversity (Torres *et al*. 2018; Johri *et al*. 2021; Barroso and Ragsdale 2025). Inbreeding via self-fertilisation also influences the strength of BGS, and although recombination rates and dominance coefficients can be rescaled to account for selfing (Nordborg 1997; Glémin and Ronfort 2013), this approach is known to be inaccurate when the rate of selfing is high enough (Kamran-Disfani and Agrawal 2014; Roze 2016). The fact that BGS is influenced by both demography and selfing suggests that the classic-BGS model may provide inaccurate estimates under these conditions. More generally, it is unclear whether this approach can be used for full parametric inference given that previous investigations often fix some of the parameters describing the distribution of deleterious fitness effects or only assess a small number of parameter combinations (Comeron 2014; Corbett-Detig *et al*. 2015; Liang *et al*. 2022).

### Overview

Here we investigate approaches for inference of BGS from genome sequence data. We use simulations to test the performance of the classic-BGS model under a range of conditions involving demographic non-equilibrium and partial-selfing. We also propose an alternative approach which aims to jointly model BGS, demography and partial-selfing, and we compare the performance of this model to classic-BGS. Our main focus is accurate estimation of the genome-wide reduction in genetic diversity due to BGS (*B̅*), but we also test the ability of these methods to estimate the distribution of deleterious fitness effects (DFE), variation in *B* along the genome (B-maps), and, where applicable, the demographic history of the population. We apply our proposed joint inference method to two species of *Capsella* with contrasting mating systems and conclude by discussing recent progress and outstanding challenges in estimation of BGS from sequence data.

## Results

### Constant population size and strong selection

We implemented the classic-BGS model as a method for estimating the reduction in genetic diversity due to BGS along the genome (*B*). Our implementation is similar to those of previous work (Comeron 2014; Elyashiv *et al*. 2016) (see Methods for details). One notable difference is that we do not use the approximation for *B* derived by Hudson and Kaplan (1995) and Nordborg *et al*. (1996), and instead use:

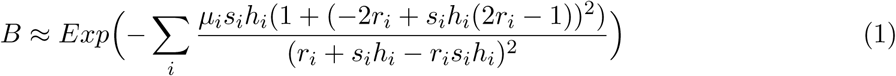

from Roze (2016), where *µ_i_* is the deleterious mutation rate at selected site *i*, *s_i_* is the reduction in fitness in a homozygote, *h_i_* is the dominance coefficient, and *r_i_* is the recombination distance to the selected site. Unlike the approximations of Hudson and Kaplan (1995) and Nordborg *et al*. (1996), Equation 1 holds under loose linkage and for unlinked mutations (*r_i_* = 1*/*2 ; see Matheson and Masel 2024 for an investigation of this issue). We assume that all selected sites share the same *µ* and *h* but that fitness effects of deleterious mutations (DFE) are gamma distributed and therefore parameterised by mean *s̅* and shape *β*. There are four free parameters in the model – *N_max_*, *µ*, *s̅*, *β* – and we assume *h* = 0.5 throughout as this parameter is not identifiable. Here *N_max_* refers to the effective population size in the absence of BGS, so that the overall reduction in genetic diversity (*B̅*) is equal to *N_e_/N_max_*. The problem of estimating *B̅*, which is our main quantity of interest, can therefore be rephrased as estimation of *N_max_*. Before fitting this model to data, we can first gain intuition about how each DFE parameter affects patterns of linked diversity by visualising analytic predictions of *B*. Figure 1 shows how varying either *µ*, *s̅* or *β* across a factor of ten, while holding the other two parameters constant, affects the predicted value of *B* given a single site under purifying selection at recombination distance *r*. Varying *µ* affects *B* equally across all values of *r*. By contrast, varying *s̅* changes the relative contributions of tightly and loosely linked sites to BGS. The shape parameter *β* determines the contribution of sites at an intermediate distance to BGS, with the overall effect being subtle. These results suggest that it will much more challenging to precisely estimate *β* than *µ* or *s̅*.

**Figure 1:**
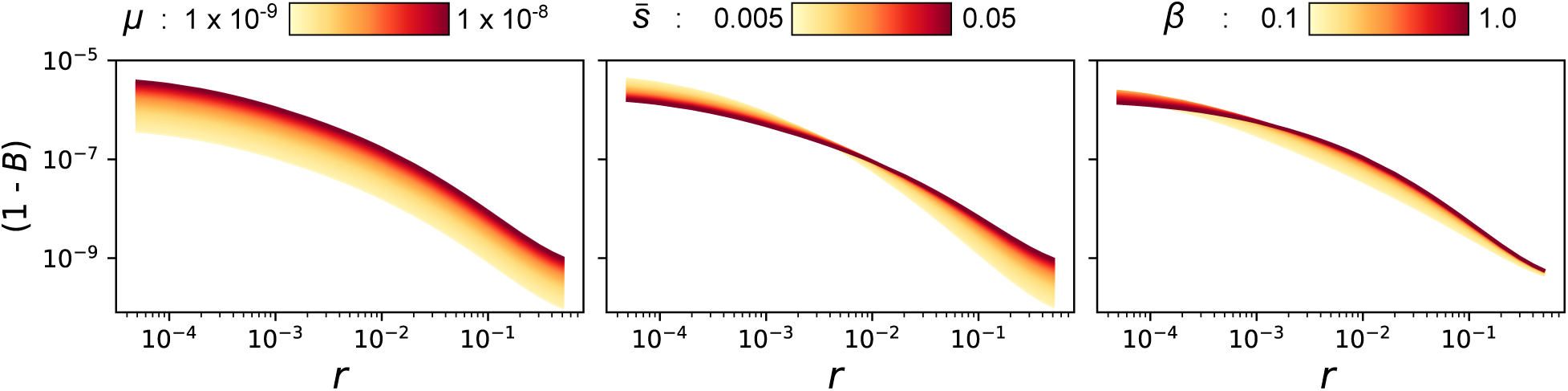
Analytic predictions for the reduction in genetic diversity from BGS. The predicted departure from neutral levels of diversity (1 − *B*, y-axis) is plotted against the recombination distance of a selected site (*r*, x-axis). Each panel shows the effect of varying a single DFE parameter across an order of magnitude, while holding the other parameters constant (*µ* = 5.5 × 10^−9^, *s̅* = 0.0275, *β* = 0.55) and assuming a coalescent *N_e_* of 10,000.

To test the performance of the classic-BGS model under favourable conditions we simulated a panmictic population of constant size where mutations within coding DNA sequence (CDS) are subject to strong purifying selection (*N* = 10,000, *µ* = 7.5 × 10^−9^, *s̅* = 0.01, *β* = 3, see Methods for more details). These simulations were conditioned on the CDS annotation and recombination map of three *Capsella rubella* chromosomes (Slotte *et al*. 2013), totaling 49 Mb in length. In these simulations BGS reduced genetic diversity within intergenic regions by 19%, i.e. *B̅* = 0.81, and this reduction varied across the genome due to variation in recombination rate and gene density (Figure S1A). We fit the classic-BGS model to these simulated data, using nucleotide diversity across 10 kb windows at input. The B-map predicted by the model is a qualitatively good fit to the B-map generated by the simulation (Figure S1A) and maximum composite likelihood (MCL) parameter estimates (*N_max_* = 9997, *µ* = 7.21 × 10^−9^, *s̅* = 0.0087, *β* = 3.08) are also a good match to the parameters of the simulation. We estimated the uncertainty in parameters by bootstrapping across the simulation replicates. While the 95% CIs are narrow for the *N_max_* parameter (9957 − 10,049), they are wider for *µ* (7.03 − 7.56 × 10^−9^) and *s̅* (0.0070 − 0.0111), and very broad for *β* (1.23 − 10.00) (Figure S1B), which is consistent with the fact that *β* has a subtle effect on *B*. These results show that the classic-BGS model can accurately estimate *N_max_*, and by extension the reduction in genetic diversity due to BGS – *B̅* – but that variation in nucleotide diversity across the genome is only weakly informative about the DFE.

Our simulations are conditioned on the recombination map of *C. rubella* (Slotte *et al*. 2013; Brazier and Glémin 2022), where recombination varies on a scale of 100 kb. We assume this map when fitting the model, yet in reality most estimated recombination maps will tend to be more coarse than the true map. To investigate this issue we repeated the analysis conditioning on maps where recombination rate is instead measured in intervals of 1 or 5 Mb. We find that coarse recombination maps lead to poorly fitting B-maps and biased parameter estimates (Figure S1). However, *N_max_* is still well estimated when using these recombination maps, albeit with more uncertainty (Figure S1). This suggests that accurate estimation of *B̅* is possible even with only partial information about recombination.

### Non-equilibrium demography

We next consider the effect of non-equilibrium demography on estimation of BGS parameters. We modified the simulation procedure to include an instantaneous five-fold growth / decline in population size at time *T* generations in the past, where *T* is equal to *N* in the most recent epoch (see Methods). Values of *N* were chosen to ensure that populations would have the same levels of nucleotide diversity at the time of sampling in the absence of BGS. Within these simulations the average reduction in diversity due to BGS was greater for the decline demography (*B̅* = 0.75) than for the growth demography (*B̅* = 0.89) (Figure 2A). This interaction between demography and BGS has been reported previously (Torres *et al*. 2018; Johri *et al*. 2021; Barroso and Ragsdale 2025). It can be explained by BGS increasing the rate of coalescence and by extension the probability that lineages coalesce in earlier epochs, so more frequently when *N* is small (see the Appendix of Johri *et al*. 2021). We fit the classic-BGS model to these simulated datasets and found that in both cases the predicted B-map provides a good fit to the simulated B-map (Figure 2A). The *N_max_* parameter (which for non-equilibrium demographies corresponds to the overall rate of coalescence in the absence of BGS) is slightly overestimated for the decline demography, but the bias is very small (∼ 1%; Figure 2B). Estimates of *s̅* and *β* have similar accuracy and precision to those from simulations with constant population size. However, we find that the deleterious mutation rate is estimated incorrectly for both non-equilibrium demographies (Figure 2B). This parameter (*µ* = 7.50×10^−9^) is underestimated under a scenario of population growth (3.96×10^−9^, (3.85−4.16)) and overestimated under population decline (10.40 × 10^−9^, (9.91 − 12.32)). This bias can be understood by considering the interaction between demography and BGS described above: the effect of BGS on nucleotide diversity is modulated by changes in population size and, because demography is missing from the model, this is accounted for by changing *µ*. So although the classic-BGS model provides meaningful estimates of *N_max_* under demographic non-equilibrium, estimates of *µ* become biased.

**Figure 2:**
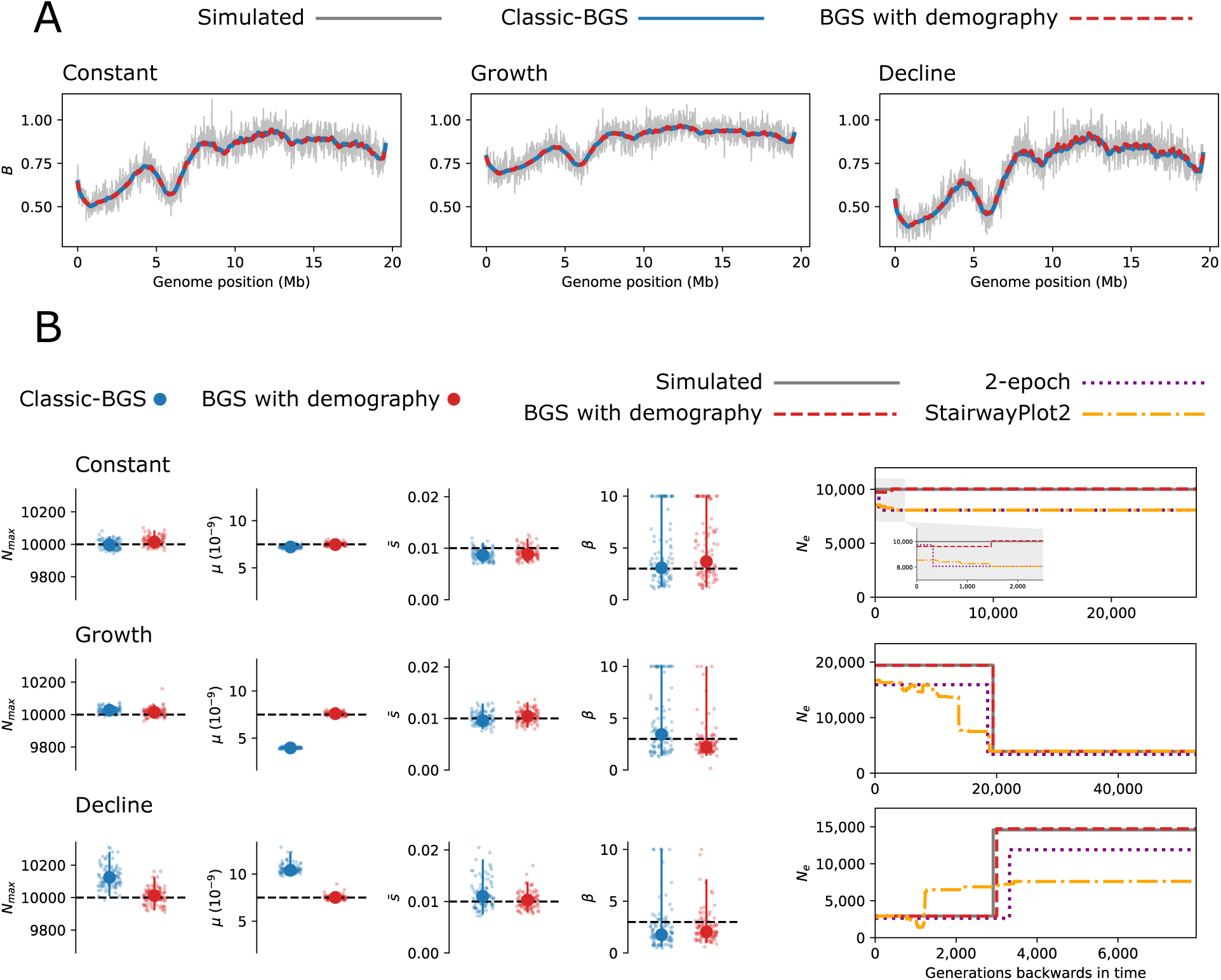
Estimation of BGS under non-equilibrium demography. **(A):** The reduction in genetic diversity from BGS along one chromosome is shown for simulations with constant population size, population growth and population decline. The grey line in each panel is the observed reduction in the simulation, whereas estimates from the classic-BGS and BGS-with-demography models are shown as blue and dashed-red lines, respectively. **(B)**: Estimates of *N_max_*, *µ*, *s̅* and *β*. Point estimates are shown as large points and bootstrap estimates are shown as small jittered points. Vertical lines show 95% CLs and dashed horizontal lines correspond to the simulated parameter values. The demographic history of each simulation scenario is shown on the right. The grey line corresponds to the number of individuals (*N*) of the simulated Wright-Fisher population. Estimates from the BGS-with-demography model are shown as a dashed-red line and the estimated history from methods assuming selective neutrality are also plotted for comparison.

Barroso and Ragsdale (2025) have also recently shown that demographic non-equilibrium leads to biased DFE inference under the classic-BGS model and have developed an elegant two-locus method to resolve this issue. Here we investigate an alternative approach for jointly estimating BGS and demography that uses information contained in site frequency spectra (SFS) along the genome. The approach is similar to the classic-BGS model with some notable differences. First, we calculate the composite likelihood of observing windowed SFS along the genome, rather than levels of nucleotide diversity. Second, we include a two-epoch demographic history and estimate *B* values separately for each epoch. Finally, we account for the transition in the rate of coalescence from *N_max_* to *BN_max_* in recent time using an approximation based on the results of Nicolaisen and Desai (2013). In summary, we model the rate of coalescence within each genomic window using demographic parameters that are shared genome-wide (*N*_0_, *N*_1_, *T*) along with epoch-specific *B* values for each window that depend on the local effect of BGS through time (*B*_0_, *B*_1_) given the local recombination rate, density of selected sites and the deleterious DFE (*µ*, *s̅*, *β*).

We fit this model of BGS-with-demography to the simulated datasets analysed in the previous section, which differ only in their demographic history (constant, growth and decline). We find that the B-maps predicted by this model are almost identical to those predicted by the classic-BGS model (Figure 2A). While the classic-BGS model failed to accurately estimate *µ* for histories of non-equilibrium demography, the new model provides accurate estimates of *µ* for all three simulated datasets (Figure 2B). The small bias in estimates of *N_max_* under the decline demography is also resolved (Figure 2B). The BGS-with-demography model estimates a two-epoch demographic history that approximately accounts for the effect of BGS on sequence variation. These estimated histories provide excellent matches to the simulated demographic parameters (Figure 2B). As a comparison, we estimated demography from the same simulated datasets using a neutral two-epoch model as well as StairwayPlot2 (Liu and Fu 2020) (which also assumes selective neutrality). We find that both of these methods generate estimates of *N_e_* that are lower than the simulated *N* values by approximately a factor of *B̅* (Figure 2B). Additionally, these methods incorrectly estimate the timing of changes in *N_e_*, especially for the decline demography. For the constant population size simulations these methods estimate a small increase in *N_e_* in recent time, corresponding to the transition between *N_max_* and *BN_max_* (Nicolaisen and Desai 2013; Figure 2B). By contrast, the model of BGS-with-demography, which aims to capture this transition in the rate of coalescence, has only a very small amount of error in recent time (*N* is underestimated by 2.8%). Although the model of BGS-with-demography that we consider is approximate in nature, these results show that it can resolve bias associated with the classic-BGS model and also provide estimates of demography that account for the effect of BGS on sequence variation.

### Weak selection

So far we have only simulated strong purifying selection where almost all deleterious mutations will be removed by natural selection. Specifically, we simulated a deleterious DFE with *s̅* = 0.01 and *β* = 3, where the large value of *β* ensures that most mutations will satisfy the condition 2*Nsh* ≫ 1. However, estimates of the DFE across a wide range of species have shown that *β* tends to be *<* 1 (Chen *et al*. 2017), suggesting that the DFE often has a considerable fraction of weakly deleterious mutations. The model of BGS-with-demography presented above can likely accommodate weak selection to some degree because the DFE is truncated at *s* = 3*/*(2*N_e_h*) within each epoch (see Methods). Truncating the DFE in this way is nonetheless an approximation and so inference could become biased if weakly deleterious mutations are sufficiently common. To investigate the effect of weak selection we repeated the simulations above while setting *β* = 0.333. This value is similar to estimates of *β* for *Drosophila melanogaster* (Campos *et al*. 2017) and implies that 17% of mutations will have fitness effects where 2*Nsh <* 1 when *s̅* = 0.01 and *N* = 10,000.

The overall reduction in genetic diversity due to BGS in these simulations was *B̅* = 0.85, 0.92, 0.81 for constant, growth and decline demographies, respectively, which are smaller reductions than observed under strong purifying selection (see previous section). We fit both the classic-BGS and BGS-with-demography models to the simulated data. The B-maps predicted by these two models are almost identical and provide a qualitatively good fit to the simulations (Figure S2A). We again find that the BGS-with-demography model generates accurate estimates of the demographic history, whereas methods that assume neutrality are biased (Figure S2B). However, we find that the DFE parameters are poorly estimated by both models (Figure S2B). This can be partly explained by the fact that when selection is weak the effective rate of deleterious mutations that contribute to BGS is determined by both *µ* and *β*, and *β* is a hard parameter to estimate when only considering the signal left by BGS at neutral sites (Figure 1). We also suspect that the piece-wise constant assumption of our model may introduce bias whenever deleterious allele frequencies are slow to equilibrate. These results again show that the DFE is challenging to estimate from this type of model. Importantly, *N_max_*, B-maps, and demographic histories are accurately estimated under the BGS-with-demography model, even when the DFE is not.

### Partial selfing

To investigate the effect of partial selfing on estimation of BGS we modified the general simulation procedure used above to include reproduction by self-fertilisation at rate *α*. We simulated a mixed-mating population (*α* = 0.4) and a high-selfing population (*α* = 0.9), both with a constant population size of *N* = 10,000 diploids and a deleterious DFE that includes the possibility of weakly selected mutations (*s̅* = 0.01, *β* = 0.333). We find that overall nucleotide diversity is reduced by a factor of 0.63 in the mixed-mating population and by a factor of 0.23 in the high-selfing population. However, these reductions include both the effects of selfing and BGS. Using knowledge of *α*, these values can be rescaled so that they only represent the effect of BGS: with *B̅* = 0.80 and *B̅* = 0.43 for mixed-mating and high-selfing, respectively. Given that *B̅* = 0.85 for a randomly-mating population with the same size and DFE (see above), this shows that the interaction between selfing and BGS is weak when populations reproduce by mixed-mating but substantial under high-selfing.

We fit the classic-BGS model to these simulated data. Estimates of *N_max_* from this model are underestimates of the simulated *N*. This is expected given that partial selfing increases the long-term rate of coalescence by a factor of (1 + *F*) (where *F* = *α/*(2 − *α*)) and the probability that a pair of lineages reach this long-term process when sampled from a diploid is only (1 − *F*). The classic-BGS model assumes random mating (i.e. *F* = 0) and so underestimates *N_max_*. The DFE parameter estimates from the classic-BGS model broadly match the true values for the mixed-mating population, albeit with a significant upwards bias in estimates of *s̅*. Estimates for the high-selfing population are strongly biased (Figure 3). Similarly, the predicted B-map is a qualitatively good fit for mixed-mating, but the effect of BGS on genetic diversity is underestimated for the high-selfing population. The classic-BGS model is therefore surprisingly robust to reproduction by mixed-mating but gives inaccurate estimates of the DFE and *B̅* under high-selfing.

**Figure 3:**
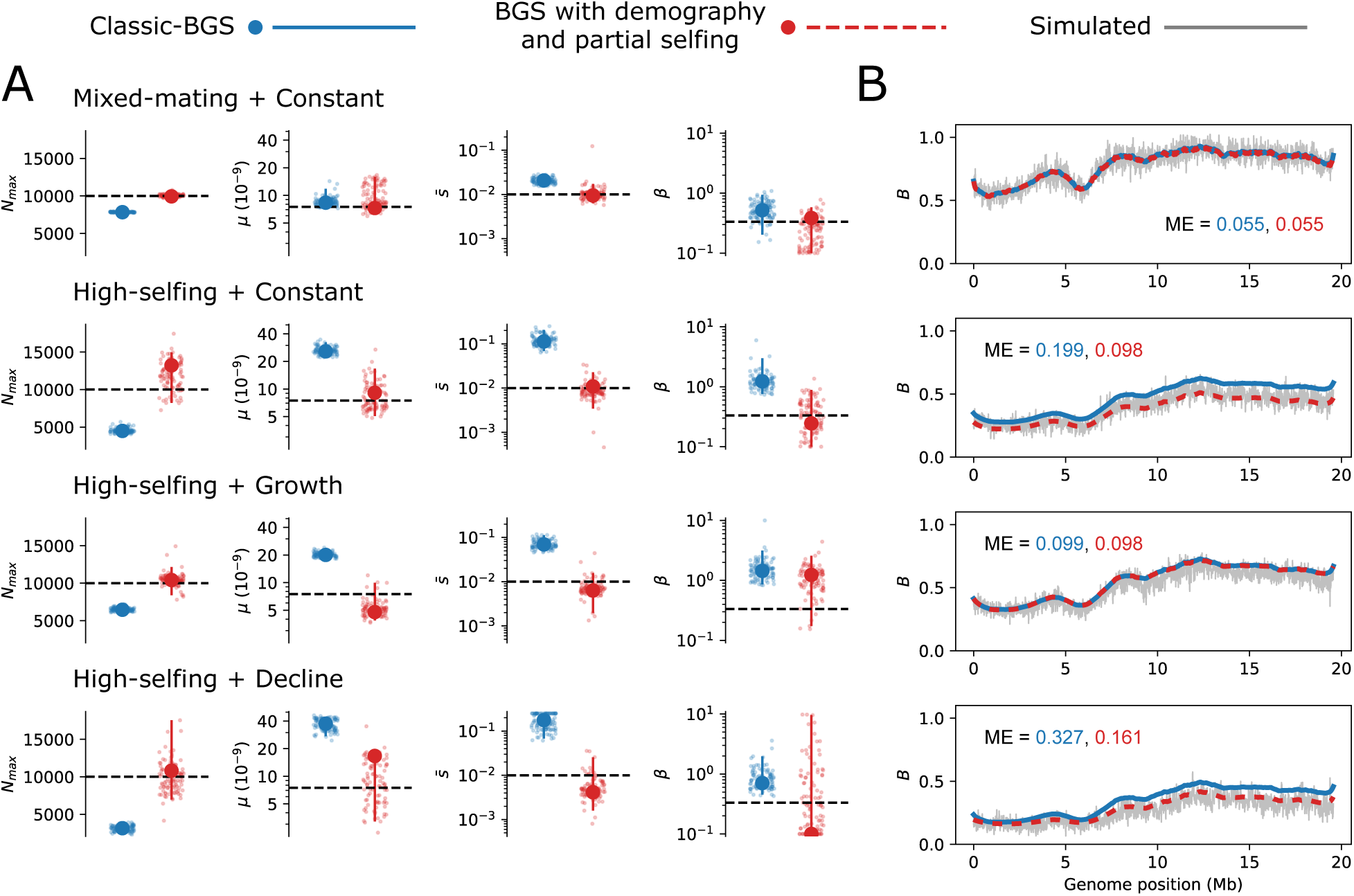
Estimation of BGS under partial selfing and non-equilibrium demography. **(A)**: Parameter estimates from the classic-BGS and BGS-with-demography model under different rates of partial selfing and demography. As in Figures S1 and 2, large points are point estimates, small jittered points are bootstrap estimates and vertical lines correspond to 95% CIs. A *log* y-axis is used for plots showing estimates of *s̅* and *β*. **(B):** The reduction in genetic diversity from BGS along one chromosome is shown for each simulated scenario. The grey line in each panel is the observed reduction in the simulation, whereas estimates from the classic-BGS and BGS-with-demography models are shown as blue and dashed-red lines, respectively. The mean error (ME) between the simulated and estimated B-maps is also shown in each panel.

The BGS-with-demography model is expected to have the same limitations when fit to data from partially selfing populations. It is, however, possible to extend the model to account for partial selfing. First, the expected *B* value for a given time epoch and genomic window can be calculated using an approximation for the effect of BGS on the rate of coalescence under partial selfing. Here we use a new approximation based on the results of Roze (2016):

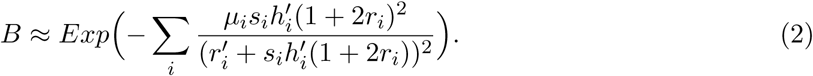

where *r*^′^ = *r*(1 − *F*) and *h*^′^ = *h*(1 − *F*) *F*. This approximation is expected to better capture the contribution of loosely linked loci to BGS than that of Nordborg (1997). Second, the expected SFS for a genomic window should reflect the increased rate of coalescence from selfing, as well as the distortion in shape due to sampling diploid individuals. This has been addressed by Blischak *et al*. (2020) and here we use their inbreeding aware implementation of *∂a∂i* to obtain the SFS for each genomic window. Together these changes allow estimation of BGS parameters conditional on *α*, which can be estimated from *F_IS_* or the SFS prior to model fitting, or left as a free parameter.

We reanalysed the data from simulations including partial selfing using this selfing-aware model of BGS-with-demography while conditioning each analysis on the value of *α* implied by *F_IS_*. Under mixed-mating, the model provides accurate parameter estimates, resolving the bias in *s̅* (Figure 3A). Under high-selfing, parameter estimates have more variance but are again approximately unbiased (Figure 3A). The predicted B-map for the high-selfing population is also a better fit than that from the classic-BGS model (Figure 3B). In principle, this model should also perform well for partially selfing populations at demographic non-equilbrium. We therefore simulated partially selfing populations with growth and decline demographies (as above) and fit both the classic and new model to these data. We find that the new model always generates a better fitting B-map, but that the improvement in mean error depends on demography (small for growth but substantial for decline). Estimates of *µ* and *s̅* from the new model are also always more accurate, although there is evidence for bias in some cases (e.g. estimates of *β* for a scenario of high-selfing with growth; Figure 3A). We performed additional simulations under high-selfing with varying *µ* and found that estimation is less accurate when the deleterious mutation rate is high (Tables S1 and S2), but that the new model consistently performs better than classic-BGS. Altogether, these results show that it is possible to obtain better estimates of the effect of BGS on genetic diversity by explicitly modeling demography and partial selfing.

While BGS leads to a small bias in the results of demographic inference under random mating (Figure 2), we expect the bias introduced under partial-selfing to be much larger. We can assess this by visualising the coalescent *N_e_* implied by the distribution of pairwise coalescence times from simulated tree sequences. Figure 4A shows that in very recent time *N_e_* is small due to coalescence from self-fertilisation in immediate ancestors of diploid individuals. Further back in time, *N_e_* reflects coalescence in the wider population, the rate of which is determined by demography, BGS and selfing. Although we simulated a piece-wise constant history, *N_e_* is not constant within epochs. In recent time this is due to the transition in coalescence rate between *N_e_* and *BN_e_* as lineages move through fitness classes. After this transition, *N_e_* tends to gradually increase again (see Discussion). The large departure between the simulated demography and realised rate of coalescence shows that demographic inference in selfing populations is challenging.

**Figure 4:**
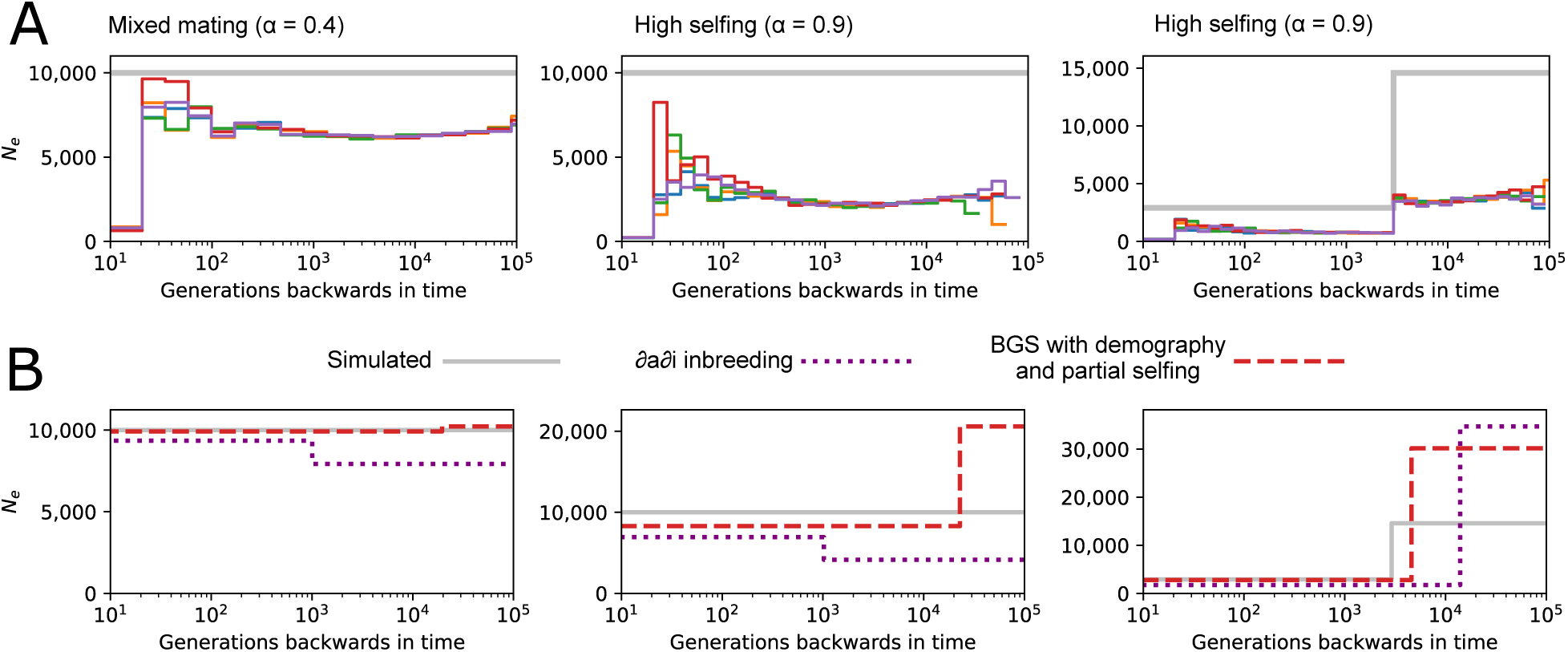
The effect of BGS and partial-selfing on effective population size (*N_e_*) through time. **(A)**: The *N_e_* implied by the distribution of pairwise coalescence from simulations with either mixed-mating (*α* = 0.4) or high-selfing (*α* = 0.9). The grey line in each plot corresponds to the simulated population size and coloured lines correspond to coalescent *N_e_* trajectories for five batches of simulation replicates. **(B)**: Point estimates of inferred demographic histories from these simulated data using either the method of Blischak *et al*. (2020) or the BGS-with-demography model extended to partial-selfing.

The two-epoch demographic histories estimated by the BGS-with-demography model are shown in Figure 4B. As a comparison we also estimated past demography using the method of Blischak *et al*. (2020), which accounts for partial selfing but not BGS. Generally, we observe that the method of Blischak *et al*. (2020) underestimates *N_e_*, estimates recent growth despite constant population sizes, and also has trouble estimating the timing of demographic changes (Figure 4B). The BGS-with-demography model performs better in these respects, but does tend to overestimate *N_e_* in deeper time when the rate of selfing is high. It is nonetheless encouraging that our approach can provide meaningful estimates of *N_max_* through time in selfing populations where the impact of BGS on patterns of sequence polymorphism is strong.

### The effect of BGS on genetic diversity in *Capsella grandiflora* and *C. orientalis*

We next investigate the role of BGS in determining levels of genetic diversity in two closely related plant species; *Capsella grandiflora*, which is a self-incompatible outcrosser, and *C. orientalis*, which reproduces predominantly by selfing. We reanalyse whole genome sequence data from Josephs *et al*. (2015) and Koenig *et al*. (2019) by jointly calling variants for 50 *C. grandiflora* and 16 *C. orientalis* samples (see Methods). Estimates of the inbreeding coefficient *F_IS_* reflect the contrasting mating systems of these species, with *F_IS_* = 0.044 in *C. grandiflora* and *F_IS_* = 0.87 in *C. orientalis*. Hereafter we assume that *C. grandiflora* mates randomly (*α* = 0) and that *C. orientalis* reproduces by partial selfing (0 *< α <* 1). Nucleotide diversity per-site (excluding CDS) is approximately fortyfold higher in *C. grandiflora* (*π* = 0.0094) than in *C. orientalis* (*π* = 0.00024), consistent with a strong effect of selection at linked sites in the predominantly selfing species.

We first fit the BGS-with-demography model to the outcrossing species *C. grandiflora*. While we have previously assumed a two-epoch demographic history and perfect polarisation of alleles when fitting this model to simulated data, here we instead assume a three-epoch history and also include an allele polarisation error parameter that we estimate under neutrality prior the model fitting (*ɛ* = 0.0536). We estimate a genome-wide reduction in diversity of *B̅* = 0.887 in *C. grandiflora*, with the B-map varying between 0.986 and 0.771 across 10 kb windows (Figure 5A). The gamma-distributed deleterious DFE for CDS is estimated with parameter values of *µ* = 3.74 × 10^−9^, *s̅* = 0.00307 and *β* = 10.0. Although this value of *β* is unrealistic, setting a realistic value gives a very similar B-map (Figure S3). The inferred demographic history is consistent with an expanding population as *N_max_* increases approximately three-fold over the last 400,000 generations (Figure 5C). Altogether, our results suggest that the effect of BGS on genetic diversity is weak in *C. grandiflora*.

**Figure 5:**
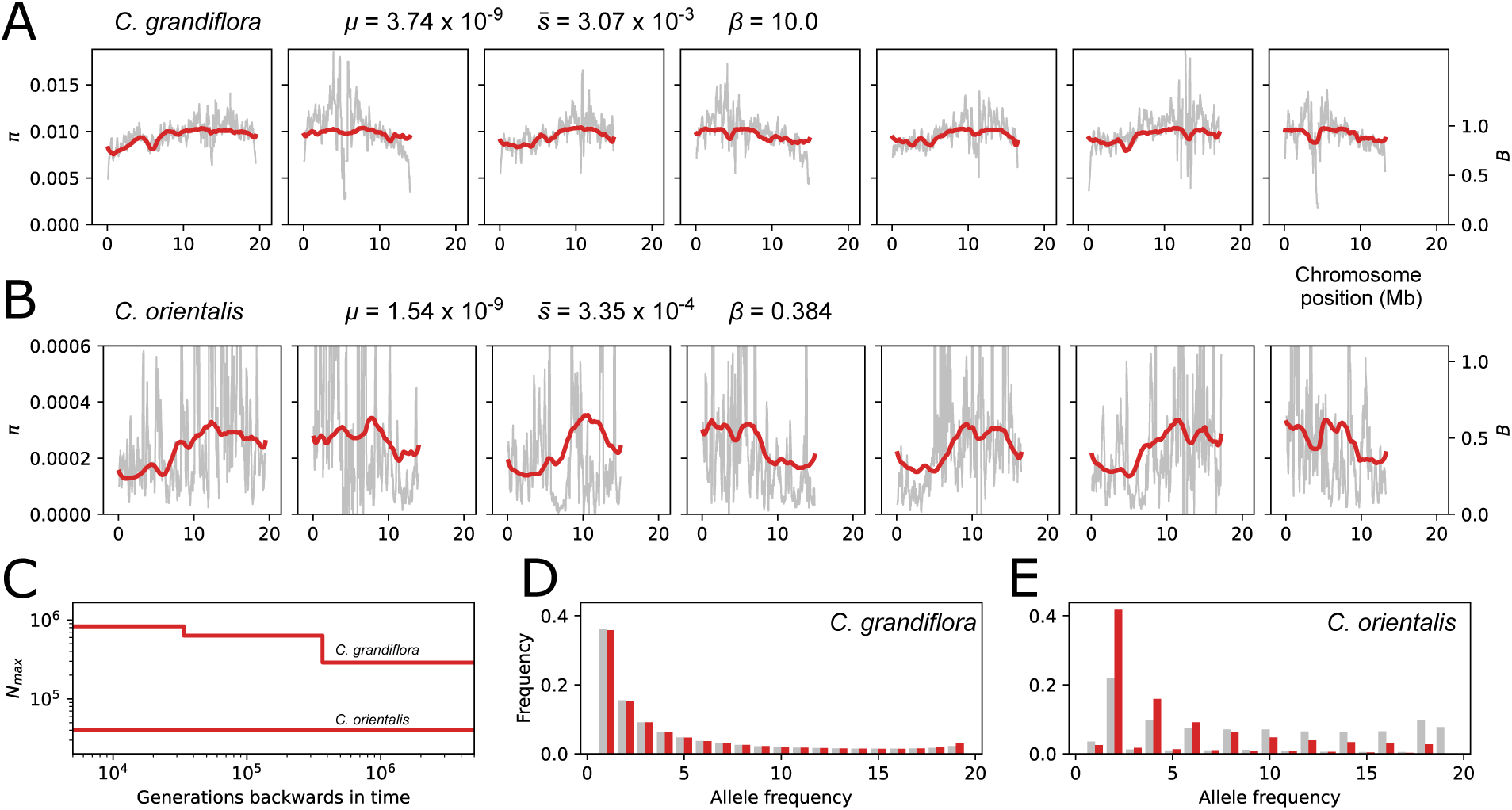
The effect of BGS on genetic diversity in *C. grandiflora* and *C. orientalis*. **(A)**: Nucleotide diversity (*π*) is plotted as a grey line across seven *C. grandiflora* chromosomes in 200 kb sliding windows. Predicted levels of *π* and *B* from BGS are shown as a red line. The estimated DFE parameters are listed above the plot. **(B)**: The same as in (A) but for *C. orientalis*. **(C)**: Estimates of *N_max_* through time for *C. grandiflora* and *C. orientalis* plotted on a log-log scale. **(D)**: The normalised unfolded SFS for *C. grandiflora* is shown as grey bars and the predicted SFS is shown as red bars. **(D)**: The same as in (B) but for *C. orientalis*.

Ideally we would fit the same model of BGS-with-demography to the predominantly selfing species *C. orientalis* while also including the effect of partial selfing. Unfortunately, the SFS for our sample of *C. orientalis* genomes is very flat (Figure 5E), which makes inference of past demography challenging. For example, fitting a neutral two-epoch demographic model to this data together with rates of polarisation error and selfing suggests a 50-fold decrease in *N_e_* at *T* = 83,383, with *ɛ* = 0.232 and *α* = 0.981. These unrealistic parameter estimates (especially *ɛ*) suggest that the shape of the SFS in *C. orientalis* may reflect other evolutionary forces that are not captured by our model (see Discussion). With this in mind, we choose to fit a simpler model of BGS-with-partial-selfing (conditional on *α* = 0.981) using only variation in *π* along the genome rather than the full SFS. Although this means we cannot include past demography in our analysis, we avoid the possibility of biasing our inference by assuming that variation in the shape of the SFS across the genome reflects BGS.

For *C. orientalis* we estimate a genome-wide reduction in diversity of *B̅* = 0.444 from BGS and a corresponding *N_max_* of 40,328 (Figure 5). The estimated DFE suggests very weak selection, with *µ* = 1.54×10^−9^, *s̅* = 3.35×10^−4^ and *β* = 0.384. Note that these DFE estimates are informed by the sojourn time of deleterious mutations, which we expect to be increased by any negative linkage-disequilrbium between selected mutations. Although our simulations suggest that this effect is small when *α* = 0.90, it is likely substantial in *C. orientalis* given *α* ≈ 0.98 and so may explain why we estimate such a weak DFE. Although we have not used the SFS when fitting this model, we can nonetheless obtain an expected SFS for this scenario of BGS-with-partial-selfing. We find that the expectation from the model, which incudes no demographic change or polarisation error, is a poor match to the data (Figure 5E), especially in comparison to the model fit for *C. grandiflora* (Figure 5D).

Our results suggest that the reduction in genetic diversity due to BGS (1 − *B̅*) is five-fold greater in *C. orientalis* than *C. grandiflora*. The fact that we simultaneously model partial selfing, the DFE and past demography means that we can also estimate the contribution of each of these processes to the difference in *B̅* between the two species. For example, while our model of *C. grandiflora* predicts *B̅* = 0.887 genome-wide, if we replace the expanding history of *C. grandiflora* with a constant population size with equivalent overall *N_max_* then we instead predict *B̅* = 0.878. This shows that demography only plays a small role in modulating the effect of BGS on genetic diversity in *C. grandiflora*, and that most of the difference in *B̅* between these species is instead due their contrasting mating systems.

## Discussion

Here we have evaluated the performance of two different methods for estimating the effect of BGS on genetic diversity: the classic-BGS model (Hudson and Kaplan 1995; Nordborg *et al*. 1996) and an SFS-based method that aims to jointly model BGS, demography and partial-selfing. Consistent with previous work, we found that demographic change (Torres *et al*. 2018; Johri *et al*. 2021; Barroso and Ragsdale 2025) and partial-selfing (Nordborg 1997; Kamran-Disfani and Agrawal 2014; Roze 2016; Burgarella *et al*. 2024) modulate the effect of BGS on genetic diversity and therefore lead to biased results when using the classic-BGS model. However, these biases mostly affect parameters describing the DFE rather than the predicted B-map. As a result, the classic-BGS model appears to be robust to both demographic change and low rates of partial-selfing if one is only interested in estimating the overall effect of BGS on genetic diversity – *B̅*. This robustness can be understood by considering the identifiability of the different parameters in the model. While the DFE parameters are estimated from patterns of variation in diversity along the genome (Figure 1), information about *N_max_* is mostly contained in regions where *B* is close to 1.0. The B-maps from simulations of randomly mating and mixed-mating populations often had considerable regions of chromosomes where *B* ≈ 1.0 (Figures 2 and 3), making estimation of *N_max_* (and *B̅*) straightforward. By contrast, B-maps from simulations of high-selfing populations tended to have values of *B* that were much lower, meaning that estimation of *N_max_* requires accurate knowledge of the DFE in order to extrapolate levels of diversity to an unobservable region that is free of BGS. Accurate inference from such populations therefore requires alternative methods, and our proposed joint inference approach appears promising in this respect (Figure 3).

Despite the robustness of the classic-BGS model, our results show that jointly modelling BGS and demography does provide several benefits. In particular, our joint inference approach resolves the bias in estimates of *µ* introduced by demographic non-equilibrium and also generates accurate inference of demography while accounting for BGS (Figure 2). These benefits are demonstrated in our analysis of *C. grandiflora*, where we could also show that a demographic history of expansion has slightly weakened the effect of BGS on genetic diversity in this species. Our approach does assume that natural selection is strong relative to drift (2*N_e_sh* ≫ 1) and that frequencies of deleterious mutations equilibrate quickly between epochs, which will not always be true when selection is weak and effective population sizes are small (Figure S2). Barroso and Ragsdale (2025) have also recently developed an approach for modelling BGS under non-equilibrium demography. Their two-locus method does not currently support joint inference (demographic histories are specified prior to BGS estimation), but it has the major advantage of explicitly modelling weak selection and deleterious allele frequencies through time. We therefore expect the method of Barroso and Ragsdale (2025) to be much more accurate than the one we propose here in regimes of weak selection. Additionally, because their approach uses polymorphism information at both selected and linked sites, we expect that estimation of DFE parameters will be more precise. While we have found that patterns of linked diversity contain limited information about the DFE (Figures 2 and 3), it will be interesting to see whether the approach of Barroso and Ragsdale (2025) will prove more powerful than DFE estimation methods that ignore the effect of selection on linked sites (Keightley and Eyre-Walker 2007).

We have shown that the classic-BGS model effectively breaks down when analysing data from populations that predominantly reproduce by selfing. By using a new approximation (Equation 2, Roze 2016) we were able to generate far more accurate estimates of *B̅* and the DFE (Figure 3). We nonetheless expect that BGS has additional effects on sequence variation in highly selfing populations that are not captured by our joint inference approach. In particular, negative linkage disequilibrium is expected to build up between deleterious mutations when sex is rare, leading to less efficient selection and deleterious mutations that segregate longer than expected given *sh* (Hill and Robertson 1966; Daigle and Johri 2025). This is evidenced by subtle underestimates of *s̅* for simulated data when *α* = 0.9 (Figure 3) and estimates of very weak selection for *C. orientalis*, where *α* ≈ 0.98. We also found that distributions of coalescence times from simulations of BGS and partial-selfing were inconsistent with a piece-wise constant demographic history (Figure 4A). While this could be due to insufficient burn-in (despite 200,000 total generations), an alternative explanation is that a regime of weak purifying selection and low effective recombination rate leads to distortions in genealogies that are not captured by a monotonic transition from *N_e_* to *BN_e_* (Seger *et al*. 2010; Strütt *et al*. 2025). For example, Khudiakova *et al*. (2024) have recently suggested that weak purifying selection results in a multiple-merger coalescent process, and although outside of the scope of this current work, it would be interesting to test whether this can explain the unusually flat SFS of *C. orientalis* (Figure 5E). We expect that our assumptions of independence between selected sites and a Kingman coalescent process may be problematic when populations are almost completely selfing. In such cases, simulation-based inference may be a better prospect for accurately estimating the effect of BGS on genetic diversity (Johri *et al*. 2021).

Several authors have suggested that BGS should be included in baseline / null models of evolution (Comeron 2014; Johri *et al*. 2020). Here we have tested whether BGS (modulated by demography and partial-selfing) can explain the large difference in genetic diversity between two closely related species – *C. grandiflora* and *C. orientalis* – or if additional demographic and selective forces are required. We estimate that the effect of BGS on genetic diversity is approximately five-fold greater in the predominant selfer *C. orientalis*, yet maximum composite likelihood estimates of *N_max_* for these species still differ by an order of magnitude (Figure 5). This suggests that much of the difference in diversity between these species is instead explained by other factors, such as fluctuations in census population size, variance in offspring number, the frequency and impact of selective sweeps (Andersen *et al*. 2012; Hartfield and Bataillon 2020), or even *de novo* mutation rates (Wang *et al*. 2023). Although it is not currently feasible to estimate the relative contribution of all these factors to levels of genetic diversity, methods that provide joint inference of a few interacting forces provide a promising approach to disentangling the determinants of genetic diversity.

## Methods

### Classic-BGS

We implemented the classic-BGS model by calculating the composite likelihood of levels of nucleotide diversity in genomic windows across the genome, given values of *N_max_*, *µ*, *s̅* and *β*. For each genomic window we calculate *B* as *B_b_B_w_*, where *B_b_* is the contribution to *B* from purifying selection elsewhere in the genome and *B_w_* is the contribution of purifying selection within that window. By integrating Equation 1 across a DFE, *B_b_* for genomic window *k* can be calculated as:

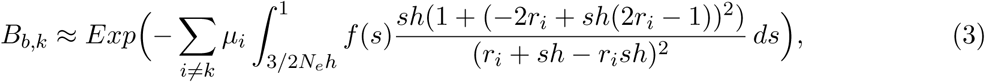

where *µ_i_* is the deleterious mutation rate in window *i* and *r_i_* is the map distance between windows *k* and *i* given by Haldane’s mapping function (Haldane 1919). When windows *k* and *i* are on different chromosomes *r_i_* = 0.5. *f* (*s*) is the probability density function of a gamma distribution parameterized by mean *s̅* and shape *β*:

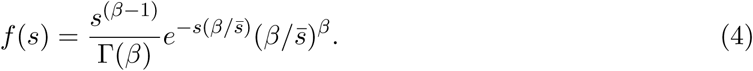

To calculate *B_w_* we assume that deleterious mutations are distributed uniformly within the window and average Equation 9 of Nordborg *et al*. (1996) across a window by integration to give:

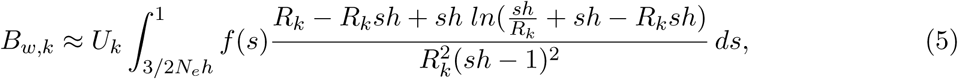

where *U_k_* and *R_k_* are the diploid deleterious mutation rate and recombination fraction in window *k*, respectively. Given *B* values for each window of the genome we calculate a log composite likelihood as:

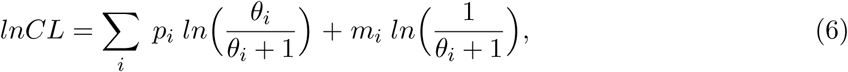

where *p_i_*and *m_i_* are counts of the number of sites in window *i* where a pair of samples are polymorphic and monomorphic, respectively, and *θ_i_*is calculated as:

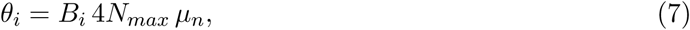

where *µ_n_* is the neutral *de novo* mutation rate per-generation. The value of *µ_n_* is set prior to model fitting and is expected to have little effect on parameter estimation as it only determines the absolute value of *N_max_* and the lower bound of the integrals in Equations 3 and 5 (as *N_e_* = *π/*(4*µ_n_*)). Even when grouping adjacent sites into windows, calculation of composite likelihoods for whole genomes can be prohibitively slow as the integrals in Equations 3 and 5 must be taken over thousands of recombination distances. To improve speed we evaluate these integrals across a grid 40 and 11 values of *r* and *R* given the parameters *s̅* and *β*, and then use linear interpolation to estimate *B_b_* and *B_w_* for each window (Comeron 2014). We perform a two-step derivative-free optimisation procedure to maximize the *lnCL*. We first perform a global search using our own implementation of the Controlled Random Search algorithm (Kaelo and Ali 2006) and then use the maximum likelihood parameter combination from this search as a starting point for a local search with the Nelder-Mead algorithm implemented in nlopt v2.7.1 (Nelder and Mead 1965; Johnson 2007).

### BGS-with-demography

We extended the classic-BGS model to jointly estimate the effects of demography and BGS on sequence variation. Demography is modelled as a piece-wise constant history where time is divided into epochs so that the rate of coalescence in epoch *t* is 1*/*(2*N_t_*) and the boundary between epoch *t* and *t* + 1 is *T_i_*. For example, in the absence of BGS and assuming a two-epoch history, the rate of coalescence through time is determined by parameters *N*_0_*, N*_1_*, T*_0_. In the presence of BGS, the coalescent history of genomic window *i* can be approximated by rescaling each *N_t_* by *B_i,t_*, where *B_i,t_* is calculated using Equations 3 and 5. One issue with these calculations is that the lower-bound of the integrals depend on the coalescent *N_e_* that results from BGS, which is unknown for individual epochs. We therefore initially assume the coalescent *N_e_* implied by genome-wide nucleotide diversity and perform iterative calculation of the B-map until *B̅_i,t_N_t_* converges.

Backwards in time, the rate of coalescence only reaches 1*/*(*B* 2*N_max_*) once all lineages in the sample are in the least loaded fitness class. Nicolaisen and Desai (2013) derived an approximation for the coalescent *N_e_* at time *t* generations in the past due to this transition:

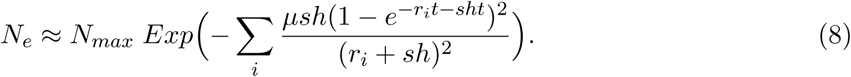

Performing calculations across values of *t* for every window in a genome would likely be prohibitively slow. We instead perform a simpler calculation using the term in Equation 8 that determines the speed of the transition, while neglecting recombination and again integrating across the DFE:

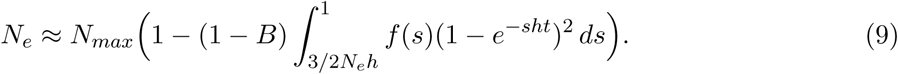

We thereby assume that the speed of the transition in coalescence rate is shared across the genome. This approximation is a drastic simplification of the coalescent process under purifying selection / BGS (Strütt *et al*. 2025). At the same time, a comparison of this approximation to those of Nicolaisen and Desai (2013) show that it is a good fit for transition dynamics in regions of low recombination (Figure S4), which is where BGS is expected to be strongest and capturing the transition in coalescence rate is most important.

For each genomic window the coalescent history is described by rescaled demographic parameters (e.g. *B*_0_*N*_0_, *B*_1_*N*_1_, *T*_0_) and a two-step transition from *N_max_* to *BN_max_* calculated by evaluating Equation 9. We then use moments v1.1.16 to calculate the expected SFS of each window given its piece-wise constant coalescent history (Jouganous *et al*. 2017). The log composite likelihood of the model given windowed SFS along the genome is calculated as:

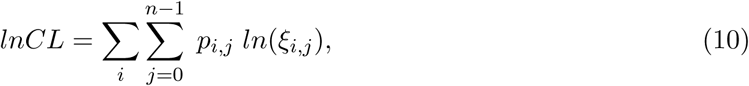

where *ξ_i,j_* is the expected frequency of sites with a derived allele count of *j* in window *i* and *p_i,j_* is the observed count of these sites. Note that the 0*^th^* entry in the SFS corresponds to monomorphic sites in the sample. We perform the same optimisation procedure as described above, but during the initial global search we round values of *B* to two decimal places to reduce the computational burden of calculating many similar expected SFS. Discretising the B-map in this way results in a substantial speed-up in computation but does also reduce the smoothness of the likelihood surface.

### BGS-with-partial-selfing

To include partial-selfing in the inference approach we integrate Equation 2 over the DFE to calculate *B_b_*. This approximation combines Nordborg’s (1997) approximation (that is obtained in the limit of weak recombination from Roze’s (2016) more general model), and Roze’s (2016) high selfing approximation. Numerical analysis indicates that the predictions obtained from this approximation are often close to the predictions from the more general model. To calculate *B_w_* and the transition in coalescence rate we rescale recombination rates and dominance coefficients by *r*(1 − *F*) and *h*(1 − *F*) + *F*, respectively, in Equations 5 and 9 (Nordborg 1997).

As above, we calculate the expected SFS for each genomic window given locally rescaled demographic parameters. Under partial-selfing coalescence rates are additionally rescaled by (1 + *F*) and we use the inbreeding aware implementation of *∂a∂i* (v2.3.3) to calculate expected spectra while accounting for the distortion introduced by sampling diploid individuals (Blischak *et al*. 2020).

Calculating B-maps and SFS along the genome under partial-selfing requires knowledge of *α*. This can be included as a free-parameter in the model, with the shape of the SFS and patterns of nucleotide diversity along the genome informing its value. We tested this approach and found that convergence of model parameters was much slower than if fixing *α* prior to model fitting. We therefore take the latter approach by calculating *F_IS_* or performing an initial estimation from the SFS under neutrality.

### Forward simulations

Forward simulations were performed with SLiM v4.2.2 (Haller *et al*. 2019; Haller and Messer 2023). Each simulation included three chromosomes that emulate chromosomes 1, 2 and 3 of the *Capsella rubella* genome. Coordinates of CDS positions corresponded to those of the genome annotation from Slotte *et al*. (2013) and recombination rates correspond to the map generated by Slotte *et al*. (2013) and later curated by Brazier and Glémin (2022). Recipes from the SLiM manual were used to model deleterious mutations within CDS, free recombination between chromosomes, instantaneous changes in population size and reproduction by partial-selfing. We simulated three different demographic scenarios: constant, growth and decline. The constant population size was *N* = 10,000, the growth parameters were *N*_0_ = 19,425, *N*_1_ = 3885, *T*_0_ = 19,425 and the decline parameters were *N*_0_ = 2919, *N*_1_ = 14,595, *T*_0_ = 2919, where *N*_0_ corresponds to the size of the population in the most recent time epoch. Simulations with random mating consisted of 12*N* generations and those with partial-selfing consisted of 12*N/*(1 + *F*) generations. We set the total number of generations to 200,000 for the simulations used to generate the results shown in Figure 4 to reduce the possibility that insufficient burn-in explained changes in coalescent *N_e_* through time. Each simulation was repeated 100 times with the output being a population-level tree sequence (Kelleher *et al*. 2018; Haller *et al*. 2019).

Tree sequences were recapitated and simplified using pySLiM v1.0.4, tskit v0.5.6 and msprime v1.3.1 (Baumdicker *et al*. 2021). Five diploid individuals were sampled per-simulation for randomly mating populations, whereas ten diploids were sampled for partial-selfing populations to ensure that estimation of *α* was not a limiting factor in our analysis. Neutral mutations were added to tree sequences while assuming a discrete genome and a per-site per-generation mutation rate of 7.5 × 10^−9^. Windowed SFS were tallied from tree sequences while removing the contribution of CDS to both polymorphic and monomorphic sites. We also outputted a VCF file for each simulation which was used to calculate *F_IS_* via the method of Weir and Cockerham (1984).

For each simulation scenario we pooled data from the 100 replicates and fit BGS models to these windowed SFS to obtain point estimates of model parameters and B-maps. We generated bootstrap replicates by randomly sampling 100 simulation replicates with replacement 100 times. These bootstrap replicates were analysed to calculate the uncertainty in parameter estimates, with upper and lower 95% CIs calculated as the 2.5 and 97.5 percentiles of parameter estimates across bootstrap replicates.

### Analysis of Capsella data

Paired-end Illumina whole genome sequence data for 50 *C. grandiflora* and 16 *C. orientalis* samples were downloaded from BioProjects PRJNA275635 and PRJEB6689, respectively (Josephs *et al*. 2015; Koenig *et al*. 2019). Mapping and variant calling was performed using a modified version of the snpArcher pipeline (Mirchandani *et al*. 2024; https://github.com/ThomasBrazier/snpArcher-dev), with the *C. rubella* genome assembly (GCF 000375325.1; Slotte *et al*. 2013) as a reference. The main modification was the inclusion of a module for calculating callable sites for each sample (≥ 10 reads but *<* 3× mean coverage) using mosdepth (Pedersen and Quinlan 2018). Callable sites were further refined by identifying and removing putative paralogous loci using ngsParalog (Linderoth 2018). The calculation for estimating excess heterozygosity used by ngsParalog was modified to account for partial-selfing. Specifically, the probability of observing a heterozygous genotype given a derived allele frequency *p* and inbreeding coefficient *F* was calculated as:

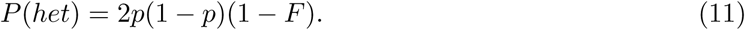

This calculation allowed us to identify paralogous loci for *C. orientalis* by calculating *F_IS_* and removing paralogs iteratively. We used the resulting intervals to define non-CDS regions where at least 10 *C. grandiflora* and 10 *C. orientalis* samples had callable sites (49 Mb in total). From these intervals we calculated unfolded SFS in 10 kb windows along the genome, with multivariate hypergeometric sampling of genotypes used to downsample spectra to 10 diploids per-species. Polarisation of alleles was performed using the genotypes of the other species, with sites showing polymorphism in both species being set to non-callable. Given the possibility of imperfect allele polarisation we estimated an error parameter (*ɛ*) for *C. grandiflora* by fitting a neutral three-epoch model to the SFS with *ɛ* as a free parameter. Under this model the expected SFS is calculated as:

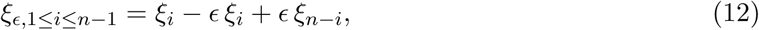

where *ξ_i_* is the expected frequency of polymorphisms with *i* derived alleles given the demographic history. The value of *ɛ* estimated under this model was assumed when fitting the BGS-with-demography model. We also estimated *ɛ* for *C. orientalis* under a neutral demographic model but found that the parameter values were unrealistic (see Results) and therefore chose to fit a model of BGS-with-partial-selfing where only levels of windowed nucleotide diversity are considered.

We used the recombination map and CDS annotation of *C. rubella* to fit BGS models, while assuming that deleterious mutations are confined to CDS. Sequence variation on chromosome 5 (KB870809.1) was excluded from the analysis because of errors in the recombination map (Brazier and Glémin 2022). Note that we included CDS positions across the entire genome in our analysis, even for chromosomes or windows with no callable sites. We assumed a *de novo* mutation rate of *µ_n_* = 7 × 10^−9^ per-site per-generation, given estimates in *Arabidopsis thaliana* (Weng *et al*. 2019), and a generation time of one year. The last two bins of the SFS (*n* − 1, *n* − 2) were masked when fitting BGS models as such polymorphisms are likely to be inflated by any reference bias. We fit BGS models using 10 replicate runs for each dataset to assess convergence. We found that the BGS-with-partial-selfing model that was fit to the *C. orientalis* data showed unsatisfactory parameter convergence across replicates and so we repeated the analysis with narrower parameter ranges. Results from repeated runs are found in Tables S3 and S4, with results from the maximum composite likelihood run reported in the main text.

## Data availability

Code for performing forward simulations with SLiM and fitting BGS models is available at https://github.com/A-J-F-Mackintosh/Binfer. Windowed spectra for the two *Capsella* species and a table of SRA accessions for samples used in the analysis can be found at the same repository.

## Acknowledgments

We would like to thank Thomas Brazier for his contributions to the modified snpArcher pipeline used in this study, Paul Blischak for advice on using *∂a∂i* with partial-selfing and Buŗcin Yıldırım for providing feedback on a previous version of this manuscript.

## References

Andersen EC, Gerke JP, Shapiro JA, Crissman JR, Ghosh R, Bloom JS, Félix MA, Kruglyak L. 2012. Chromosome-scale selective sweeps shape *Caenorhabditis elegans* genomic diversity. Nature genetics. 44:285–290.

Barroso GV, Ragsdale AP. 2025. A model for background selection in non-equilibrium populations. bioRxiv. pp. 2025–02.

Barry P, Broquet T, Gagnaire PA. 2022. Age-specific survivorship and fecundity shape genetic diversity in marine fishes. Evolution letters. 6:46–62.

Baumdicker F, Bisschop G, Goldstein D, Gower G, Ragsdale AP, Tsambos G, Zhu S, Eldon B, Ellerman EC, Galloway JG et al. 2021. Efficient ancestry and mutation simulation with msprime 1.0. Genetics. 220. iyab229.

Blischak PD, Barker MS, Gutenkunst RN. 2020. Inferring the demographic history of inbred species from genome-wide SNP frequency data. Molecular biology and evolution. 37:2124–2136.

Brazier T, Glémin S. 2022. Diversity and determinants of recombination landscapes in flowering plants. PLoS Genetics. 18:e1010141.

Buffalo V. 2021. Quantifying the relationship between genetic diversity and population size suggests natural selection cannot explain Lewontin’s paradox. Elife. 10:e67509.

Burgarella C, Brémaud MF, Von Hirschheydt G, Viader V, Ardisson M, Santoni S, Ranwez V, de Navascués M, David J, Glémin S. 2024. Mating systems and recombination landscape strongly shape genetic diversity and selection in wheat relatives. Evolution Letters. 8:866–880.

Campos JL, Zhao L, Charlesworth B. 2017. Estimating the parameters of background selection and selective sweeps in *Drosophila* in the presence of gene conversion. Proceedings of the National Academy of Sciences. 114:E4762–E4771.

Charlesworth B. 2012. The effects of deleterious mutations on evolution at linked sites. Genetics. 190:5–22.

Charlesworth B, Jensen JD. 2022. How can we resolve Lewontin’s paradox? Genome biology and evolution. 14:evac096.

Charlesworth B, Morgan M, Charlesworth D. 1993. The effect of deleterious mutations on neutral molecular variation. Genetics. 134:1289–1303.

Chen J, Glémin S, Lascoux M. 2017. Genetic diversity and the efficacy of purifying selection across plant and animal species. Molecular biology and evolution. 34:1417–1428.

Comeron JM. 2014. Background selection as baseline for nucleotide variation across the *Drosophila* genome. PLoS Genetics. 10:e1004434.

Coop G. 2016. Does linked selection explain the narrow range of genetic diversity across species? BioRxiv. p. 042598.

Corbett-Detig RB, Hartl DL, Sackton TB. 2015. Natural selection constrains neutral diversity across a wide range of species. PLoS biology. 13:e1002112.

Daigle A, Johri P. 2025. Hill-Robertson interference may bias the inference of fitness effects of new mutations in highly selfing species. Evolution. 79:342–363.

Elyashiv E, Sattath S, Hu TT, Strutsovsky A, McVicker G, Andolfatto P, Coop G, Sella G. 2016. A genomic map of the effects of linked selection in *Drosophila*. PLoS genetics. 12:e1006130.

Glémin S, Ronfort J. 2013. Adaptation and maladaptation in selfing and outcrossing species: new mutations versus standing variation. Evolution. 67:225–240.

Haldane JB. 1919. The combination of linkage values, and the calculation of distances between the loci of linked factors., In:, Routledge. pp. 385–395.

Haller BC, Galloway J, Kelleher J, Messer PW, Ralph PL. 2019. Tree-sequence recording in SLiM opens new horizons for forward-time simulation of whole genomes. Molecular ecology resources. 19:552–566.

Haller BC, Messer PW. 2023. SLiM 4: multispecies eco-evolutionary modeling. The American Naturalist. 201:E127–E139.

Hartfield M, Bataillon T. 2020. Selective sweeps under dominance and inbreeding. G3: Genes, Genomes, Genetics. 10:1063–1075.

Hill WG, Robertson A. 1966. The effect of linkage on limits to artificial selection. Genetics Research. 8:269–294.

Hudson RR, Kaplan NL. 1995. Deleterious background selection with recombination. Genetics. 141:1605–1617.

Johnson SG. 2007. The NLopt nonlinear-optimization package. https://github.com/stevengj/nlopt.

Johri P, Charlesworth B, Jensen JD. 2020. Toward an evolutionarily appropriate null model: jointly inferring demography and purifying selection. Genetics. 215:173–192.

Johri P, Riall K, Becher H, Excoffier L, Charlesworth B, Jensen JD. 2021. The impact of purifying and background selection on the inference of population history: problems and prospects. Molecular biology and evolution. 38:2986–3003.

Josephs EB, Lee YW, Stinchcombe JR, Wright SI. 2015. Association mapping reveals the role of purifying selection in the maintenance of genomic variation in gene expression. Proceedings of the National Academy of Sciences. 112:15390–15395.

Jouganous J, Long W, Ragsdale AP, Gravel S. 2017. Inferring the joint demographic history of multiple populations: beyond the diffusion approximation. Genetics. 206:1549–1567.

Kaelo P, Ali MM. 2006. Some variants of the controlled random search algorithm for global optimization. Journal of Optimization Theory and Applications. 130:253–264.

Kamran-Disfani A, Agrawal A. 2014. Selfing, adaptation and background selection in finite populations. Journal of evolutionary biology. 27:1360–1371.

Keightley PD, Eyre-Walker A. 2007. Joint inference of the distribution of fitness effects of deleterious mutations and population demography based on nucleotide polymorphism frequencies. Genetics. 177:2251–2261.

Kelleher J, Thornton KR, Ashander J, Ralph PL. 2018. Efficient pedigree recording for fast population genetics simulation. PLoS computational biology. 14:e1006581.

Khudiakova KA, Boenkost F, Tourniaire J. 2024. Genealogies under purifying selection. bioRxiv. pp. 2024–10.

Koenig D, Hagmann J, Li R, Bemm F, Slotte T, Neuffer B, Wright SI, Weigel D. 2019. Long-term balancing selection drives evolution of immunity genes in *Capsella*. Elife. 8:e43606.

Leffler EM, Bullaughey K, Matute DR, Meyer WK, Ségurel L, Venkat A, Andolfatto P, Przeworski M. 2012. Revisiting an old riddle: What determines genetic diversity levels within species? PLOS Biology. 10:1–9.

Lewontin RC. 1974. The genetic basis of evolutionary change..

Liang YY, Shi Y, Yuan S, Zhou BF, Chen XY, An QQ, Ingvarsson PK, Plomion C, Wang B. 2022. Linked selection shapes the landscape of genomic variation in three oak species. New Phytologist. 233:555–568.

Linderoth T. 2018. Identifying population histories, adaptive genes, and genetic duplication from population-scale next generation sequencing. University of California, Berkeley.

Liu X, Fu YX. 2020. Stairway Plot 2: demographic history inference with folded SNP frequency spectra. Genome biology. 21:280.

Mackintosh A, Laetsch DR, Hayward A, Charlesworth B, Waterfall M, Vila R, Lohse K. 2019. The determinants of genetic diversity in butterflies. Nature communications. 10:3466.

Matheson J, Masel J. 2024. Background Selection From Unlinked Sites Causes Nonindependent Evolution of Deleterious Mutations. Genome Biology and Evolution. 16:evae050.

McVicker G, Gordon D, Davis C, Green P. 2009. Widespread genomic signatures of natural selection in hominid evolution. PLoS genetics. 5:e1000471.

Mirchandani CD, Shultz AJ, Thomas GW, Smith SJ, Baylis M, Arnold B, Corbett-Detig R, Enbody E, Sackton TB. 2024. A fast, reproducible, high-throughput variant calling workflow for population genomics. Molecular Biology and Evolution. 41:msad270.

Murphy DA, Elyashiv E, Amster G, Sella G. 2022. Broad-scale variation in human genetic diversity levels is predicted by purifying selection on coding and non-coding elements. Elife. 12:e76065.

Nelder JA, Mead R. 1965. A simplex method for function minimization. The Computer Journal. 7:308–313.

Nicolaisen LE, Desai MM. 2013. Distortions in genealogies due to purifying selection and recombination. Genetics. 195:221–230.

Nordborg M. 1997. Structured coalescent processes on different time scales. Genetics. 146:1501– 1514.

Nordborg M, Charlesworth B, Charlesworth D. 1996. The effect of recombination on background selection. Genetics Research. 67:159–174.

Pedersen BS, Quinlan AR. 2018. Mosdepth: quick coverage calculation for genomes and exomes. Bioinformatics. 34:867–868.

Pope NS, Singh A, Childers AK, Kapheim KM, Evans JD, López-Uribe MM. 2023. The expansion of agriculture has shaped the recent evolutionary history of a specialized squash pollinator. Proceedings of the National Academy of Sciences. 120:e2208116120.

Romiguier J, Gayral P, Ballenghien M, Bernard A, Cahais V, Chenuil A, Chiari Y, Dernat R, Duret L, Faivre N et al. 2014. Comparative population genomics in animals uncovers the determinants of genetic diversity. Nature. 515:261–263.

Roze D. 2016. Background selection in partially selfing populations. Genetics. 203:937–957.

Seger J, Smith WA, Perry JJ, Hunn J, Kaliszewska ZA, Sala LL, Pozzi L, Rowntree VJ, Adler FR. 2010. Gene genealogies strongly distorted by weakly interfering mutations in constant environments. Genetics. 184:529–545.

Slotte T, Hazzouri KM, Àgren JA, Koenig D, Maumus F, Guo YL, Steige K, Platts AE, Escobar JS, Newman LK et al. 2013. The *Capsella rubella* genome and the genomic consequences of rapid mating system evolution. Nature genetics. 45:831–835.

Smith JM, Haigh J. 1974. The hitch-hiking effect of a favourable gene. Genetics Research. 23:23–35.

Stoffel M, Humble E, Paijmans A, Acevedo-Whitehouse K, Chilvers B, Dickerson B, Galimberti F, Gemmell N, Goldsworthy S, Nichols H et al. 2018. Demographic histories and genetic diversity across pinnipeds are shaped by human exploitation, ecology and life-history. Nature Communications. 9:4836.

Strütt S, Excoffier L, Peischl S. 2025. A generalized structured coalescent for purifying selection without recombination. Genetics. p. iyaf013.

Torres R, Szpiech ZA, Hernandez RD. 2018. Human demographic history has amplified the effects of background selection across the genome. PLoS genetics. 14:e1007387.

Wang Y, McNeil P, Abdulazeez R, Pascual M, Johnston SE, Keightley PD, Obbard DJ. 2023. Variation in mutation, recombination, and transposition rates in *Drosophila melanogaster* and *Drosophila simulans*. Genome Research. 33:587–598.

Weir BS, Cockerham CC. 1984. Estimating F-statistics for the analysis of population structure. evolution. pp. 1358–1370.

Weng ML, Becker C, Hildebrandt J, Neumann M, Rutter MT, Shaw RG, Weigel D, Fenster CB. 2019. Fine-grained analysis of spontaneous mutation spectrum and frequency in *Arabidopsis thaliana*. Genetics. 211:703–714.

