## Supplementary Materials for "Estimating the reduction in genetic diversity from background selection under non-equilibrium demography and partial selfing"

### Supplementary Figures

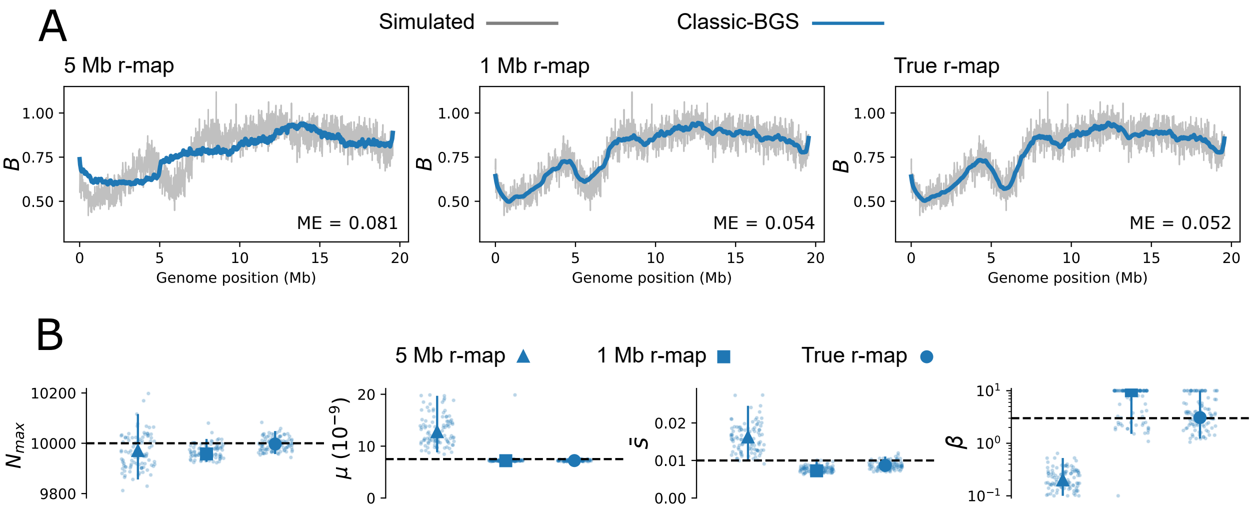


Figure S1: B-maps and parameter estimates from the classic-BGS model under varying recombination map resolution. **(A):** The simulated reduction in nucleotide diversity ($B$) generated by BGS along one simulated chromosome is shown in grey, with the estimated B-map from the classic-BGS model shown as a blue line. The mean error (ME) between the simulated and predicted B-map is shown in the bottom-right corner of each panel. The panels vary in the resolution of the recombination map that was used for model fitting. **(B):** Estimates of $N_{max}$, $\mu$, $\bar{s}$ and $\beta$ from the classic-BGS model. Point estimates are shown as large points, with shape corresponding to the resolution of the recombination map. Bootstrap estimates are shown as small jittered points and 95% CLs are shown as vertical lines. Dashed horizontal lines in each panel correspond to the simulated parameter values.

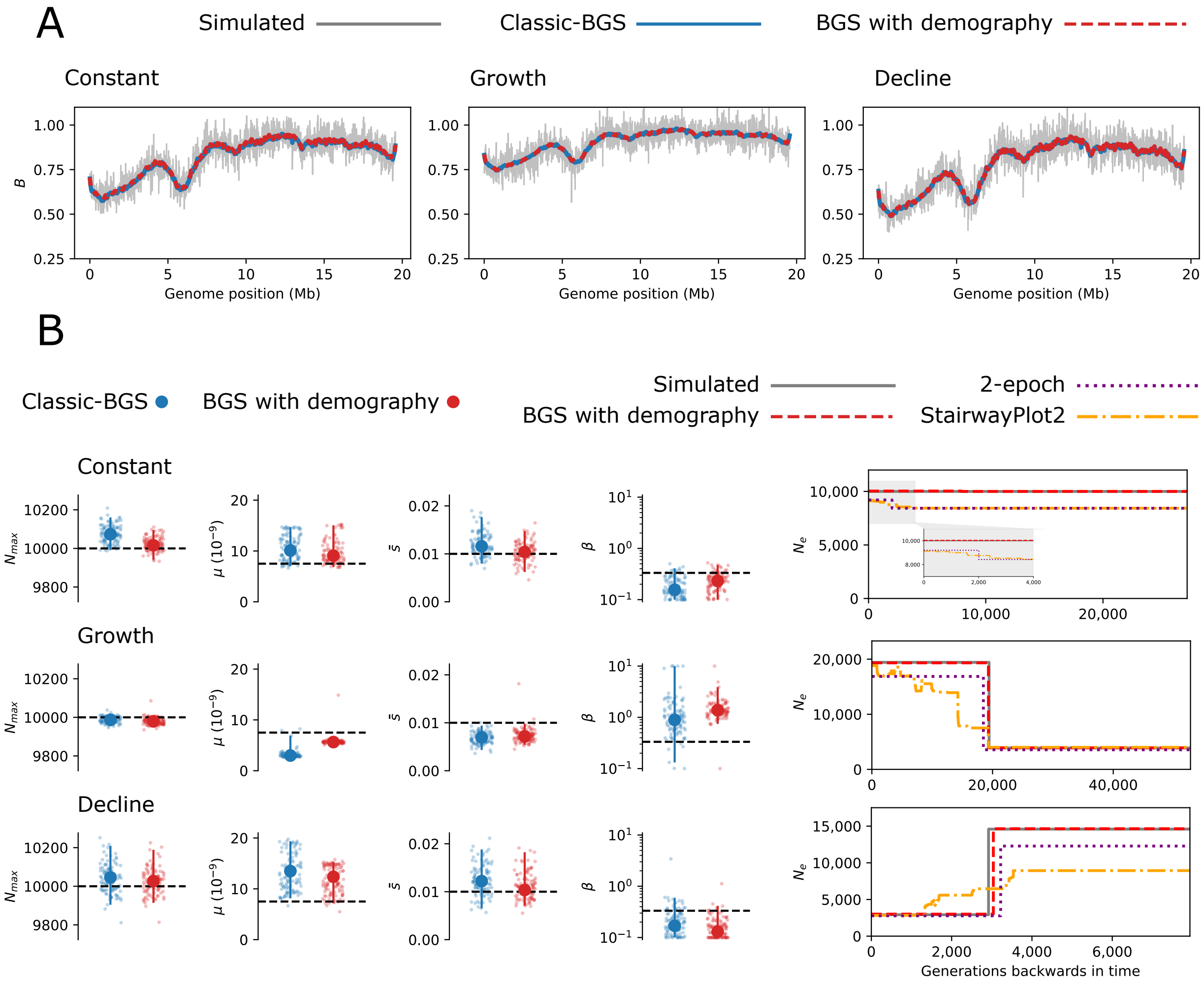


Figure S2: Estimation of BGS under non-equilibrium demography and weak selection. **(A):** The reduction in genetic diversity from BGS along one chromosome is shown for simulations with constant population size, population growth and population decline. The grey line in each panel is the observed reduction in the simulation, whereas estimates from the classic-BGS and BGS-with-demography models are shown as blue and dashed-red lines, respectively. **(B)**: Parameter estimates from the classic-BGS and BGS-with-demography model under each demographic scenario (see Figure S1). The demographic history of each simulation scenario is shown on the right. The grey line corresponds to the number of individuals ($N$) of the simulated Wright-Fisher population. Estimates from the BGS-with-demography model are shown as a dashed-red line and the estimated history from methods assuming selective neutrality are also plotted for comparison.

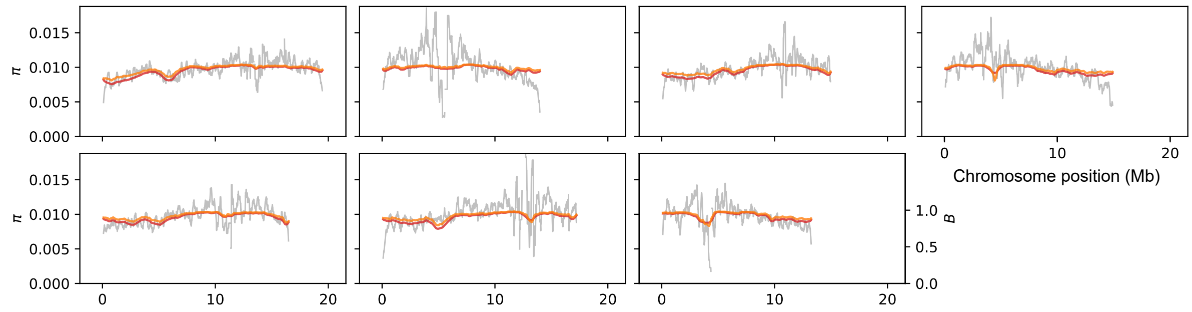


Figure S3: Levels of nucleotide diversity ($\pi$) and $B$ in *C. grandiflora*. The grey line corresponds to observed levels of $\pi$ in sliding $200$ kb windows. The red line shows the prediction from the BGS-with-demography model, which includes the parameter value $\beta = 10.0$. The orange line shows the prediction when setting $\beta = 0.2$ but retaining all other parameter values.

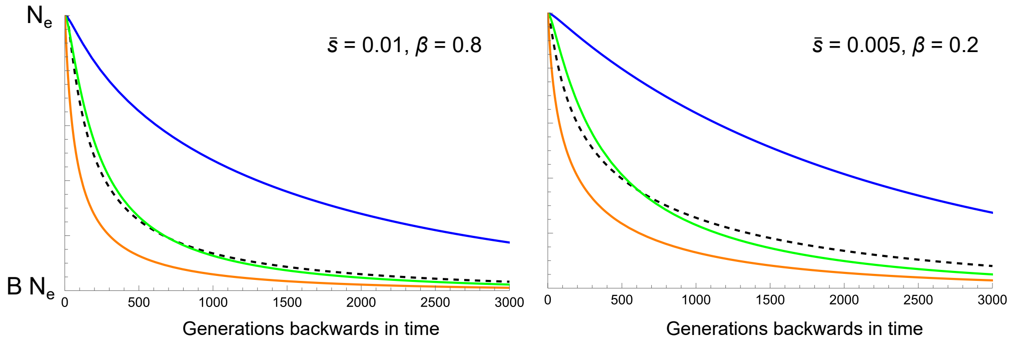


Figure S4: Approximations for the transition in coalescent $N_e$ under BGS. The coloured lines correspond to predictions from Equation 8 (Nicolaisen and Desai 2013) when integrating across a genetic map of length $50$ (orange), $0.5$ (green) or $0$ cM (blue). The prediction from Equation 9, which is used in the BGS-with-demography model, is shown as a black-dashed line. The two plots shows predictions for different DFE shapes, with weaker selection leading to slower transitions.

### Supplementary Tables

Table S1: Maximum composite likelihood parameter estimates from fitting models of BGS to data from a simulated population with high-selfing ($\alpha = 0.9$) and a low rate of deleterious mutation ($\mu = 3.75 \times 10^{-9}$). The first row shows the parameters used in the simulation, whereas the lower rows show estimates from the BGS-with-demography and partial-selfing model and the classic-BGS model. For the classic-BGS model the estimate of $N_0$ (i.e. $N_{max}$) in brackets has been rescaled to account for partial-selfing. Note that $\alpha$ is estimate prior to model fitting from $F_{IS}$ and $\bar{B}$ is a prediction from the model rather than a parameter.

| | $N_0$ | $N_1$ | $T_0$ | $\mu$ | $\bar{s}$ | $\beta$ | $\alpha$ | $\bar{B}$ |
|---|---|---|---|---|---|---|---|---|
| Simulated | 10,000 | - | - | $3.75 \times 10^{-9}$ | 0.0100 | 0.333 | 0.900 | 0.647 |
| Demography + selfing | 9138 | 20,022 | 34,069 | $4.24 \times 10^{-9}$ | 0.0051 | 0.232 | 0.902 | 0.653 |
| Classic | 4855 (9251) | - | - | $13.42 \times 10^{-9}$ | 0.1044 | 0.689 | - | 0.702 |

Table S2: Maximum composite likelihood parameter estimates from fitting models of BGS to data from a simulated population with high-selfing ($\alpha = 0.9$) and a high rate of deleterious mutation ($\mu = 1.5 \times 10^{-8}$). See the caption of Table S1 for more details.

| | $N_0$ | $N_1$ | $T_0$ | $\mu$ | $\bar{s}$ | $\beta$ | $\alpha$ | $\bar{B}$ |
|---|---|---|---|---|---|---|---|---|
| Simulated | 10,000 | - | - | $1.50 \times 10^{-8}$ | 0.0100 | 0.333 | 0.900 | 0.222 |
| Demography + selfing | 7859 | 6914 | 473 | $2.21 \times 10^{-8}$ | 0.0095 | 0.136 | 0.903 | 0.307 |
| Classic | 2645 (5041) | - | - | $3.15 \times 10^{-8}$ | 0.1133 | 1.444 | - | 0.443 |

Table S3: Maximum composite likelihood parameter estimates from fitting the BGS-with-demography model to *C. grandiflora* data ten times. Repeated runs are sorted by descending $lnCL$.

| $N_0$ | $N_1$ | $N_2$ | $T_0$ | $T_1$ | $\mu$ | $\bar{s}$ | $\beta$ | $lnCL$ |
|---|---|---|---|---|---|---|---|---|
| 830,505 | 634,878 | 289,169 | 33,873 | 367,875 | $3.738\times10^{-9}$ | 0.003067 | 10.000 | −5,586,958.52 |
| 831,344 | 635,049 | 289,210 | 33,672 | 367,699 | $3.738\times10^{-9}$ | 0.003067 | 10.000 | −5,586,958.52 |
| 828,258 | 634,673 | 289,156 | 34,369 | 368,027 | $3.739\times10^{-9}$ | 0.003070 | 10.000 | −5,586,958.52 |
| 828,426 | 634,919 | 289,205 | 34,221 | 367,686 | $3.738\times10^{-9}$ | 0.003068 | 9.937 | −5,586,958.54 |
| 831,002 | 635,251 | 289,306 | 33,793 | 367,701 | $3.747\times10^{-9}$ | 0.003143 | 9.985 | −5,586,958.56 |
| 832,633 | 635,477 | 289,408 | 33,603 | 367,777 | $3.758\times10^{-9}$ | 0.003222 | 9.999 | −5,586,958.66 |
| 821,980 | 632,641 | 288,524 | 35,543 | 368,195 | $3.785\times10^{-9}$ | 0.003312 | 5.669 | −5,586,959.17 |
| 830,080 | 634,986 | 289,259 | 34,050 | 368,013 | $3.749\times10^{-9}$ | 0.003212 | 6.221 | −5,586,960.18 |
| 792,806 | 625,091 | 288,681 | 48,837 | 374,977 | $3.783\times10^{-9}$ | 0.003500 | 4.499 | −5,586,962.81 |
| 684,340 | 278,077 | 2,791,517 | 342,698 | 3,101,951 | $3.476\times10^{-9}$ | 0.003145 | 10.000 | −5,586,995.75 |

Table S4: Maximum composite likelihood parameter estimates from fitting the BGS-with-partial-selfing model to *C. orientalis* data ten times. All runs assume $\alpha = 0.981$ and are sorted by descending $lnCL$.

| $N_{max}$ | $\mu$ | $\bar{s}$ | $\beta$ | $lnCL$ |
|---|---|---|---|---|
| 40,329 | $1.540 \times 10^{-9}$ | $3.349 \times 10^{-4}$ | 0.3840 | −53,220.87 |
| 40,781 | $1.634 \times 10^{-9}$ | $3.339 \times 10^{-4}$ | 0.3427 | −53,220.88 |
| 41,457 | $1.934 \times 10^{-9}$ | $3.106 \times 10^{-4}$ | 0.2562 | −53,220.90 |
| 39,699 | $1.067 \times 10^{-9}$ | $4.239 \times 10^{-4}$ | 0.8536 | −53,220.93 |
| 41,963 | $3.437 \times 10^{-9}$ | $1.924 \times 10^{-4}$ | 0.1163 | −53,220.97 |
| 42,382 | $3.349 \times 10^{-9}$ | $2.065 \times 10^{-4}$ | 0.1184 | −53,220.98 |
| 42,098 | $3.667 \times 10^{-9}$ | $1.836 \times 10^{-4}$ | 0.1069 | −53,220.99 |
| 42,101 | $3.694 \times 10^{-9}$ | $1.824 \times 10^{-4}$ | 0.1060 | −53,220.99 |
| 38,748 | $9.532 \times 10^{-10}$ | $4.104 \times 10^{-4}$ | 1.2500 | −53,221.00 |
| 104,952 | $3.565 \times 10^{-9}$ | $9.991 \times 10^{-3}$ | 0.1001 | −53,225.77 |
